# Dysregulation of Neuropilin-2 Expression in Inhibitory Neurons Impairs Hippocampal Circuit Development and Enhances Risk for Autism-Related Behaviors and Seizures

**DOI:** 10.1101/2024.02.05.578976

**Authors:** Deepak Subramanian, Carol Eisenberg, Andrew Huang, Jiyeon Baek, Haniya Naveed, Samiksha Komatireddy, Michael W. Shiflett, Tracy S. Tran, Vijayalakshmi Santhakumar

## Abstract

Dysregulation of development, migration, and function of interneurons, collectively termed interneuronopathies, have been proposed as a shared mechanism for autism spectrum disorders (ASDs) and childhood epilepsy. Neuropilin-2 (Nrp2), a candidate ASD gene, is a critical regulator of interneuron migration from the median ganglionic eminence (MGE) to the pallium, including the hippocampus. While clinical studies have identified Nrp2 polymorphisms in patients with ASD, whether selective dysregulation of Nrp2-dependent interneuron migration contributes to pathogenesis of ASD and enhances the risk for seizures has not been evaluated. We tested the hypothesis that the lack of Nrp2 in MGE-derived interneuron precursors disrupts the excitation/inhibition balance in hippocampal circuits, thus predisposing the network to seizures and behavioral patterns associated with ASD. Embryonic deletion of Nrp2 during the developmental period for migration of MGE derived interneuron precursors (iCKO) significantly reduced parvalbumin, neuropeptide Y, and somatostatin positive neurons in the hippocampal CA1. Consequently, when compared to controls, the frequency of inhibitory synaptic currents in CA1 pyramidal cells was reduced while frequency of excitatory synaptic currents was increased in iCKO mice. Although passive and active membrane properties of CA1 pyramidal cells were unchanged, iCKO mice showed enhanced susceptibility to chemically evoked seizures. Moreover, iCKO mice exhibited selective behavioral deficits in both preference for social novelty and goal-directed learning, which are consistent with ASD-like phenotype. Together, our findings show that disruption of developmental Nrp2 regulation of interneuron circuit establishment, produces ASD-like behaviors and enhanced risk for epilepsy. These results support the developmental interneuronopathy hypothesis of ASD epilepsy comorbidity.

## Introduction

Autism spectrum disorders (ASD) and epilepsy are highly comorbid conditions that are proposed to share common pathophysiological mechanisms. Anomalies in the development, migration, and circuit function of interneurons, collectively termed interneuronopathies, are closely associated with ASD and epilepsy ^1–4^. In particular, the development of seizures and behavioral impairments observed in several ASD subtypes are associated with altered function of inhibitory neurons. GABAergic interneurons play a pivotal role in the organization and function of the hippocampus, a brain region frequently associated with ASD and implicated in epilepsy ^5,6^. Disruptions in the establishment of cortical interneuron circuits can compromise hippocampal network function, potentially serving as a common factor contributing to the observed high comorbidity between ASD and epilepsy ^7–12^. Here we report that developmental dysregulation of a classical guidance cue receptor Neuropilin-2 (Nrp2), specifically in developing interneurons, compromises hippocampal circuit function and predisposes the network to seizures and behavioral deficits consistent with ASD.

During embryonic development, the class 3 secreted semaphorins and their obligate binding receptors, the neuropilins, are key regulators of neuronal migration, axonal guidance, dendritic morphology, and synaptic specificity of various cell types ^13–15^. Notably, Nrp2 is a strong candidate ASD gene on SFARI ranking system (Score 2) with two reported de-novo polymorphisms in Nrp2 gene in patients meeting diagnostic criteria for ASD using DSM-IV/DSM-V, Childhood Autism Rating Scale, and Autism Behavior Checklist ^16,17^. In excitatory neurons, Nrp2 expression contributes to pruning of synapses, spines and axons, whereas, in inhibitory neurons, Nrp2 regulates migration of interneurons from the medial ganglionic eminence (MGE) to the pallium, including the hippocampus ^18–20^. During embryonic development, the expression of Nrp2 in interneuron progenitors is tightly regulated by the transcription factor Nkx2.1 ^21^, and allows for migration of parvalbumin (PV+), somatostatin (SOM+) and neuropeptide-Y (NPY+) expressing interneurons to the cortex and hippocampus ^22^. We previously found that global constitutive knockout of Nrp2 leads to loss of hippocampal interneurons, produces behavioral phenotypes consistent with ASD, and increases seizure susceptibility ^23,24^. However, whether selective deletion of Nrp2 in interneurons alone and during their time of migration to the cortex and hippocampus, could result in ASD like behaviors and enhance risk for seizures is currently not known. We hypothesize that loss of Nrp2 during embryonic ages (E12.5-13.5) specifically in interneuron precursors derived from the MGE will result in fewer inhibitory neurons in cortical circuits and impair cognitive behaviors. Focusing on the hippocampus, we predict that the ensuing altered inhibitory and related circuit plasticity will disrupt hippocampal function and increase seizure susceptibility. To test these hypotheses, we deleted Nrp2 in MGE-derived interneuron precursors by crossing the Nrp2 flox conditional mouse with the Nkx2.1-CreERT2 driver line, where cre recombinase expression is induced by tamoxifen administered to pregnant dams on E12.5 and E13.5. Tamoxifen was administered at embryonic ages (E12.5-13.5) to activate cre recombinase expression during interneuron migration timepoints while minimizing activation during later neural processes ^25–29^. The resulting genotype *Nrp2^f/f^;Nkx2.1-Cre^+^* (iCKO) animals and littermate controls were used to examine hippocampal CA1 circuit functions and evaluated whether the lack of Nrp2 in interneuron precursors enhances the risk for development of seizures and behavioral deficits consistent with ASD.

## Materials and Methods

### Animals

All experiments were performed in accordance with IACUC protocols approved by Rutgers University, Newark, NJ, and the University of California at Riverside, CA and in keeping with the ARRIVE guidelines. The *Nrp2* floxed mouse^30^, which contains an IRES-GFP-polyA sequence inserted immediately downstream of the 3’ loxP in the targeting construct that allows the expression of GFP following cre recombination in all conditional mutant (−/−) neurons, was crossed with the *Nkx2.1-CreERT2* mouse (stock #014552, The Jackson Laboratory), to selectively target MGE-derived interneuron progenitors during embryonic development ^21^ (Supplementary Fig.1). Cre recombinase was induced by administering tamoxifen (5 mg, by oral gavage) to the pregnant dams at E12.5 and E13.5, during the peak timeline for developmental interneuron migration (Fig.1A and Supplementary Fig.1). Deletion of *Nrp2* in the progeny (*Nrp2^f/f^;Nkx2.1Cre^+^)* resulting in inhibitory conditional knockout (iCKO) mice was verified by visualization of GFP expression in the hippocampus in adult mice (Fig. 1B-C). While Nkx2.1-CreERT gene excision efficiency was not quantified in this study, previous studies^31^ report high excision efficiency. All appropriate littermates, such as Cre negative *Nrp2^+/f^;Cre^−^ or Nrp2^f/f^;Cre^−^* and *Nrp2^+/+^;Cre^+^*, were used as controls. In immunostaining and behavioral studies, data from *Nrp2^+/f^;Cre^+^*were not statistically different from Cre negative controls or *Nrp2^+/+^;Cre^+^* and were pooled for analysis. Mice used in these studies were backcrossed for at least 10 generations to the *C57BL/6NTac* background strain. *Nrp2* genotypes were confirmed by using polymerase chain reaction (PCR).

**Figure 1.**
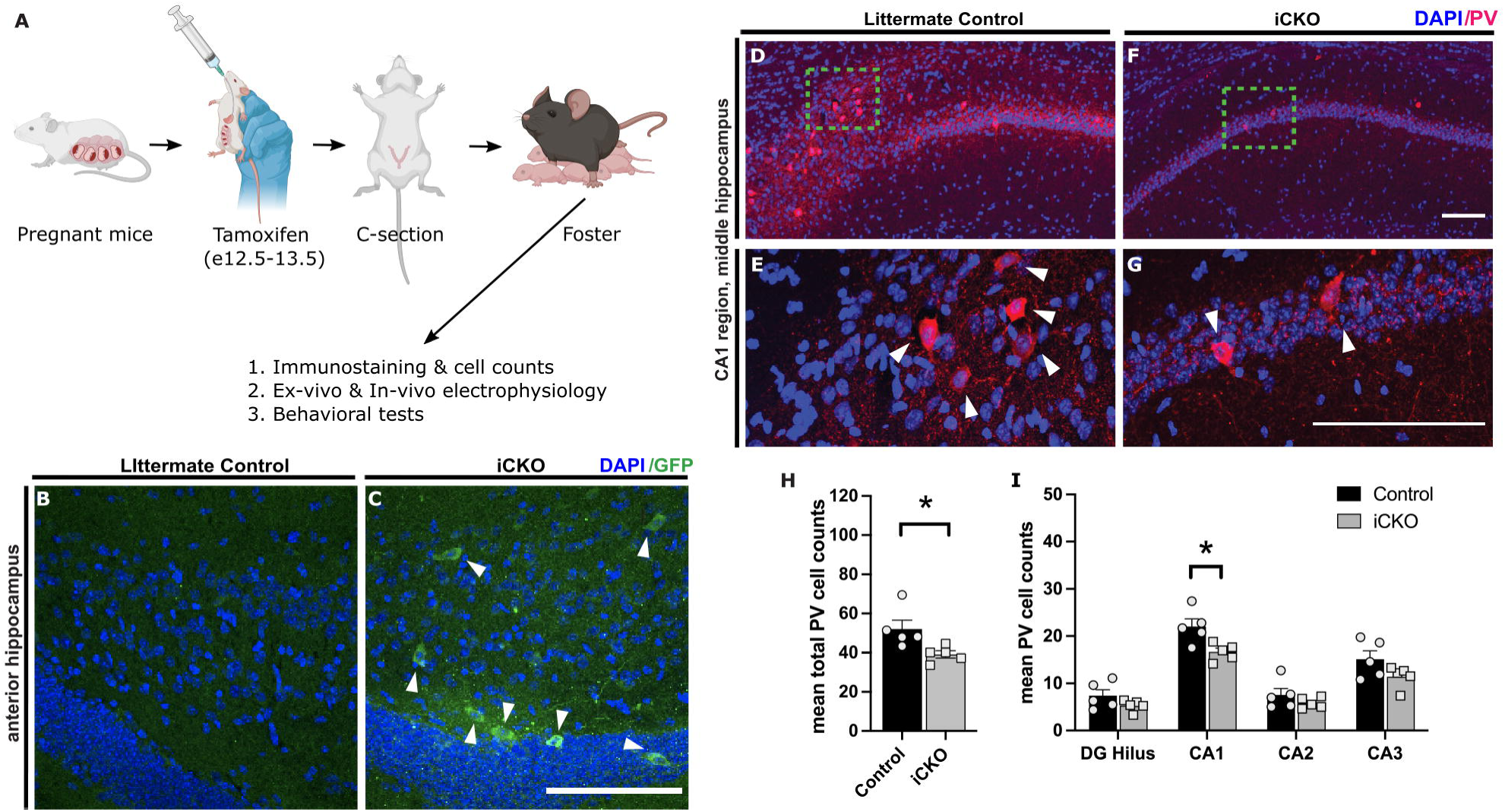
Developmental deletion of *Nrp2* leads to reduced numbers of parvalbumin (PV) expressing neurons in the hippocampus. A) Timeline adopted for excising Nrp2 in Nkx2.1 expressing neurons at E12.5 and E13.5. Tamoxifen administered at E12.5 and E13.5 by oral gavage, pups were delivered by C-section then housed with a foster mom. B, C) Brain sections immuno-labeled with anti-GFP (green) in the anterior DG region demonstrate excision of Nrp2 (mutant cells). D-G) Immuno-labeled control (D, E) and iCKO (F, G) brain sections, respectively, with anti-PV (red) and DAPI (blue). E, G) High magnification images of green boxes in D and F, respectively. H) Quantification of total mean PV+ hippocampal cells. I) Quantification of the average number of PV+ cells by hippocampal region. n=5 animals/genotype. Error bars are ± SEM; two-way ANOVA, post-hoc Bonferroni for multiple comparisons: \**p=*0.0132 CA1 region, \**p=*0.0302 unpaired *t-*test overall hippocampus. Significantly fewer number of PV+ neurons found in hippocampus of iCKO compared to control mice. All scale bars: 100 μm.

### Immunostaining and cell count

Immunostaining, cell counts, and photo-documentation were obtained from serial sections 200 µm apart across the entire hippocampus including dorsal and ventral sections (Supplementary Fig. 2) as detailed previously in Eisenberg et al 2021 ^24^ and described in detail in Supplementary Methods. Subregions were defined by The Mouse Brain in Stereotaxic Coordinates, 3rd edition ^32^.

### Ex vivo physiology

Whole-cell patch clamp recordings of CA1 pyramidal cells (CA1 PCs) were obtained from horizontal hippocampal slices (350µm) of littermate controls (*Nrp2^+/f^;Cre^−^ or Nrp2^f/f^;Cre^−^)* and iCKO mice (*Nrp2^f/f^;Nkx2.1Cre^+^*; n = 3 animals/group, 12 months old). Slice preparation and recording methods are as detailed previously ^24,33^. Voltage and current clamp recordings were obtained using MultiClamp 700B amplifiers, digitized at 10kHz using DigiData 1440A or DigiData 1550B and recorded using pClamp10 software (Molecular Devices, Sunnyvale, CA). Active and passive properties were recorded in current clamp from a holding potential of −70mV using K-Gluconate based internal solution. Voltage clamp recording from CA1 PCs held at −70mV and 0mV were used to isolate glutamatergic and GABAergic synaptic inputs, respectively, using a Cesium-based internal solution. Action potential independent miniature currents were isolated by tetrodotoxin (TTX, 1μM). Intrinsic properties were analyzed using pClamp software 10.7 (Molecular Devices, Sunnyvale, CA) and synaptic currents were detected using template search feature in Easy Electrophysiology (version 2.6). Data from cell in which access resistance changes over 20% during recordings and recordings with access resistance above 25 MΩ were excluded from analysis.

### In vivo electrophysiology

Eleven mice (5 *Nrp2^+/f^;Cre^−^ or Nrp2^f/f^;Cre^−^*littermate controls and *6 Nrp2^f/f^;NkxCre^+^*; average age of 9.12 ± 1.80 months, eight males and three females) were surgically implanted a tungsten wire depth electrode (50 μm, California Fine Wire company) in the CA1 subfield (AP:2 mm, ML: 1.5 mm, DV: 1.2mm from bregma) and a cortical screw electrode. Two additional screw electrodes (Invivo1, Roanoke, VA) on the contralateral hemisphere served as ground and reference. After 3–5 days of recovery, mice were connected to a tethered video-EEG monitoring system. Signals were sampled at 10 kHz, amplified (preamplifier: 8202-SE3, gain – x100, Pinnacle Technologies, Lawrence, KS), digitized (Powerlabs16/35, AD Instruments, Colorado Springs, CO), and recorded using LabChart 8.0 (AD instruments). Following 30 minutes of baseline recordings, mice received a single high dose of KA (20 mg/kg) to evoke seizures ^34,35^. Seizures were defined as rhythmic activity exceeding a threshold of mean baseline amplitude + 2.5 standard deviation for >5 s(12). Artifacts due to electrical, exploratory, and grooming behavior were identified and removed manually from the raw EEG’s before analysis. Latency to electrographic seizure, seizure duration, and mortality were quantified. Seizures were scored by a blinded investigator based on a modified Racine scale. Seizure severity scores in the first 30 min were averaged over 5 min epochs.

### Behavior testing

The behavior methods are fully described in the supplemental methods and in previous publications ^23^

### Social novelty test

We assessed social behavior in a three-chambered arena (Fig 6A). In the first phase, one chamber contained a caged mouse, the opposite chamber contained an empty cage, and the test mouse was able to explore both chambers. During the test for social novelty, which was conducted immediately following the first phase, one chamber contained the familiar mouse used previously, and the opposite chamber contained a novel mouse. In both phases, the test mouse explored the arena for 5 minutes. Two observers blind to experimental conditions assessed time spent sniffing the novel and familiar mouse from video footage. Testing order was block randomized across genotypes.

### Novel object recognition test

The test mouse explored two identical objects in an open field arena for 10 minutes. After a 30-min retention interval, the mouse reentered the arena, which contained an object they previously encountered (familiar object) and a novel object. Two observers blind to experimental conditions assessed time spent sniffing the novel and familiar objects from video footage. Testing order was block randomized across genotypes.

### Instrumental goal directed behavior test

We tested mice on an instrumental goal-directed learning task. Food-restricted mice pressed levers to obtain food pellets. Responses on one lever delivered chocolate-flavored pellets, whereas responses on the opposite lever delivered grain pellets. Mice underwent 36 instrumental training sessions, two sessions per day, with each lever trained in separate sessions. Mice then underwent a selective-satiety outcome devaluation procedure, in which they had free access to one flavor of food pellets in their home cage for 1 hour. In the choice test that followed, both levers were available, and mice could respond for 5 minutes, with no pellets delivered during the test. After 4 retraining sessions, we repeated the test with the opposite outcome devalued prior to the choice test. Lever-outcome assignments were randomized across genotypes.

### Statistical analysis

Unpaired *t*-tests, two-way ANOVA and post hoc Bonferroni’s, Sidak’s or Tukey’s multiple comparison correction were used to compare cell count and behavioral data. Normality was assessed for behavioral data using Q-Q plots. Homogeneity of variance was assessed using Levene’s test and Mauchly’s sphericity test for within-subject data. We found no violations of normality or variance homogeneity. Kolmogorov-Smirnov test for cumulative distributions was used to evaluate differences in distribution of synaptic parameters. Unpaired students *t*-test was used to compare differences in intrinsic properties, latency to convulsive seizures and the total time spent in seizures. Mantel-Cox test was used for survival analysis. All statistical tests were conducted in GraphPad Prism.

The difference between two dependent means (matched pairs) was used to determine the sample size requirement of behavioral tests using G*Power 3.1 software. A sample size requirement of three to seven animals for cell counts, over 2 cells/animal from a minimum of 3 mice for slice physiology and five to eight animals for behavior testing was estimated by using 80% power and an effect size found in previous work and literature. Exclusion criteria of three standard deviations from the mean was pre-established for behavior tests. One mutant mouse performed over three standard deviations lower than the mean and was therefore eliminated from behavior analysis. Significance was set to *p* < 0.05. Data are shown as mean ± SEM or median and interquartile range (IQR), as appropriate.

## Results

### Developmental deletion of *Nrp2* in interneuron progenitors reduces hippocampal interneuron numbers

Previously, Nrp2 was shown to be expressed in MGE derived interneuron precursors and is critical for cortical and hippocampal migration of interneurons ^20,36^. To examine the impact of Nrp2 expression in inhibitory neuron precursors on hippocampal circuit formation and function, we generated the *Nrp2^f/f^;Nkx2.1-CreERT2^+^*(iCKO), where cre recombinase expression is induced by feeding the pregnant dams with tamoxifen on E12.5 and E13.5 (Fig. 1A, Supplementary Fig. 1). Induction of cre-recombinase was confirmed by GFP expression in the hippocampus in a subset of sections from iCKO mice, as the *Nrp2* flox contained an IRES-GFP-pA sequence that will be shifted in frame to be transcribed upon cre recombination, but not in littermate controls (Fig.1B-C).

To determine the effects of Nrp2 deletion on the distribution of different MGE derived inhibitory neuron subtypes in hippocampal subfields, we quantified the population of PV+, SOM+ and NPY+ expressing interneurons in iCKO mice. We focused on the CA1 subfield, a region closely associated with storing social memories and pathologically implicated in ASD ^37,38^. Soma targeting PV+ interneurons are estimated to account for 21% of MGE-derived interneurons ^39^. Selective deletion of Nrp2 led to a 32.91% decrease in hippocampal PV+ interneurons population compared to control mice (Fig. 1D-I, PV neurons / section, littermate control: 52.06±4.56; iCKO: 39.17±1.81; n=5 mice, averaged over 10 sections in each, p=0.03; *t*_(8)_=2.629 unpaired *t-*test). In the CA1 region, we observed a 24.54% decrease in PV+ neuron population (mean cell count control: 22.04±1.61; iCKO: 16.63±0.81 in n=5 mice each p=0.01; t_(32)_=3.176 by two-way ANOVA with Bonferroni multiple comparison correction). We did not observe a difference in PV+ neurons in other hippocampal subfields including dentate gyrus (DG), CA2 and CA3.

Dendrite targeting SOM+ neurons are important in regulating dendritic inputs and MGE-derived interneurons account for approximately 60% of SOM+ neuron population in the hippocampus^39^. Following deletion of Nrp2, we observed a 37.12% decrease in overall hippocampal SOM+ population (Fig. 2A-F; cell count, control: 84.03±7.88; iCKO: 52.83±7.50, *p*=0.0426; *t*_(6)_=2.565 unpaired *t*-test). In the CA1 we observed a substantial 46.41% decrease in SOM+ neurons (cell count: control: 36.90±3.97; iCKO: 19.78±3.14 in n=5 mice each *p*=0.0004; *t*_(24)_=4.659 by two-way ANOVA with Bonferroni multiple comparison correction). Interestingly, there was an apparent reduction in SOM+ cells in the DG, but the difference did not reach statistical significance (*p*=0.385). We did not observe any differences in SOM+ cells in CA2 and CA3 subfields of the hippocampus.

**Figure 2.**
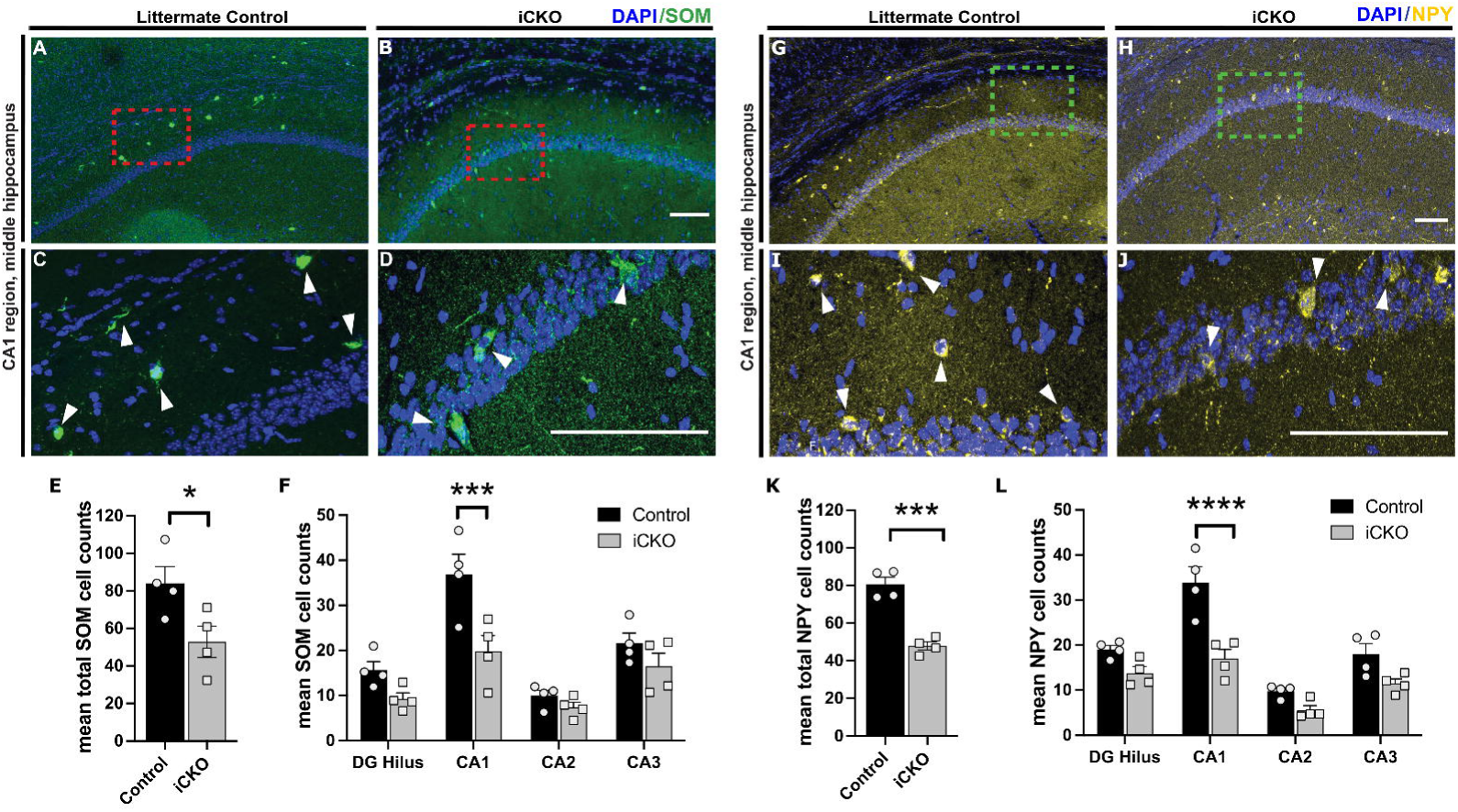
Deletion of *Nrp2* at an embryonic stage results in reduced numbers of somatostatin (SOM) and neuropeptide Y (NPY) expressing neurons in the CA1 region of the hippocampus. A-D) Immuno-labeled of littermate control (A, C,) and iCKO (B, D) brain sections, respectively, with anti-SOM (green) and DAPI (blue). High magnification images (C, D) of area in red boxes in A, B, respectively. Scale bars: 100 μm. E) Quantification of total mean SOM+ hippocampal cells. Error bars are ± SEM; unpaired *t-*test: \**p=*0.0426. Significantly fewer number of SOM+ neurons found in hippocampus of iCKO compared to control mice. F) Quantification of the average number of SOM+ cells by hippocampal region. Significantly lower SOM+ cell density found in CA1 region of iCKO mice compared to littermate controls; \*\*\**p=*0.0004. n=4 animals/genotype. G-J) Immuno-labeled of littermate control (G, I) and iCKO (H, J) brain sections, respectively, with anti-NPY (yellow) and DAPI (blue). High magnification images (I, J) of area in green boxes in G, H, respectively. Scale bars: 100 μm. K) Quantification of total mean NPY+ hippocampal cells. Error bars are ± SEM; two-way ANOVA, post-hoc Bonferroni for multiple comparisons: \*\*\**p*=0.0003. Significantly fewer number of NPY+ neurons found in hippocampus of iCKO compared to control mice. L) Quantification of the average number of NPY+ cells by hippocampal region. Significantly lower NPY+ cell density found in CA1 region of iCKO mice compared to littermate controls; \*\*\*\**p<*0.0001. n=4 animals/genotype.

We next quantified the population of NPY+ expressing interneurons which contribute to shaping synchronized activity in hippocampal circuits ^40,41^. We observed a 40.7% decrease in total NPY+ expressing interneurons in the hippocampus. (Fig. 2G-L; mean cell count: controls: 80.63±3.30; iCKO: 47.83±1.93, n=5 each *p*=0.0003; *t*_(6)_=7.674 unpaired *t*-test). In the CA1 subfield, we observed a 49.02% decrease in NPY+ neurons (mean cell count: controls: 33.90±3.10; iCKO: 17.01±1.81, *t*_(24)_=6.519; two-way ANOVA with Bonferroni multiple comparison correction *p*<0.0001). In other subfields including DG, CA2 and CA3, we observed a trend towards a decrease in NPY+ neurons (*p*=0.2135, *p*=0.4776 and *p*=0.0755 respectively) which did not reach statistical significance.

Together, our results show a significant decrease in PV+, SOM+ and NPY+ interneuron population in the CA1 following developmental deletion of Nrp2 in MGE-interneuron precursors. This reduction in interneuron populations supporting perisomatic and dendritic feedback inhibition is likely to have functional consequences at a neuronal and network level which we examined further.

### Developmental *Nrp2* deletion in interneurons alters inhibitory and excitatory synaptic transmission in CA1

Altered excitation / inhibition balance is considered a fundamental pathophysiological mechanism affecting cortical and hippocampal function in ASD ^7,8^. Decrease in interneuron populations in CA1 can alter basal inhibitory control of CA1 pyramidal cells (PCs). We examined whether action potential dependent spontaneous inhibitory currents and action potential independent miniature inhibitory synaptic inputs to CA1 were affected in iCKO mice. Voltage clamp recordings of spontaneous inhibitory postsynaptic currents (sIPSCs) in CA1 PCs revealed a significant increase in interevent intervals (IEI), indicating a decrease frequency in iCKO mice (Fig.3A,B, sIPSC interevent intervals in ms, controls: 115.3 ± 4.57, median: 71.40, IQR: 35.0 - 150.9, *n* =8 cells from 3 mice; iCKO: 133.6 ± 4.82, median: 87.65, IQR: 42.2 – 166.9, n =9 cells from 3 mice; *p* = 0.0037 by Kolmogorov-Smirnov test, Cohen’s D: 0.13). However, iCKO mice showed an increase in sIPSC amplitude compared to control mice (Fig.3B boxed inset; in pA: controls: 17.56 ± 1.086, median: 15.51, IQR: 12.46 - 21.23, iCKO: 22.29 ± 1.059, median: 20.30, IQR 16.04 – 25.51; *n*=8 vs 9 cells from 3 mice; *p*<0.0001 by Kolmogorov-Smirnov test, Cohen’s D: 0.50). While reduction in sIPSC frequency could potentially arise due reduced interneuron population in the CA1, increase in sIPSC amplitude suggests potential compensatory increase in iCKO mice ^42^. Further examination of action potential independent miniature currents (mIPSC) also revealed a significant increase in interevent intervals (decrease in frequency) in iCKO mice indicating a potential decrease in inhibitory neuron synapses on to CA1 PCs or release probability (Fig. 3C-D; mIPSC IEI in ms, controls: 88.79 ± 3.04, median: 68.45, IQR: 33.53 – 113.1, *n* =8 cells from 3 mice; iCKO : 126.5 ± 4.87, median: 85.40, IQR: 45.13 – 163.9, n = 7 cells from 3 mice; *p*<0.0001 by Kolmogorov-Smirnov test, Cohen’s D: 0.37). Unlike sIPSC amplitude, there was a decrease in mIPSC amplitude in iCKO mice (Fig. 3D boxed inset; in pA: control: 17 ± 0.25, median: 15.45, IQR 11.88 – 20.41, iCKO: 18.05 ± 0.29, median: 16.65, IQR 12.94-21.28, *p*<0.001 by Kolmogorov-Smirnov test, Cohen’s D: 0.14). Together our data show a significant reduction in spontaneous and miniature inhibitory inputs to CA1 PCs in iCKO mice.

**Figure 3:**
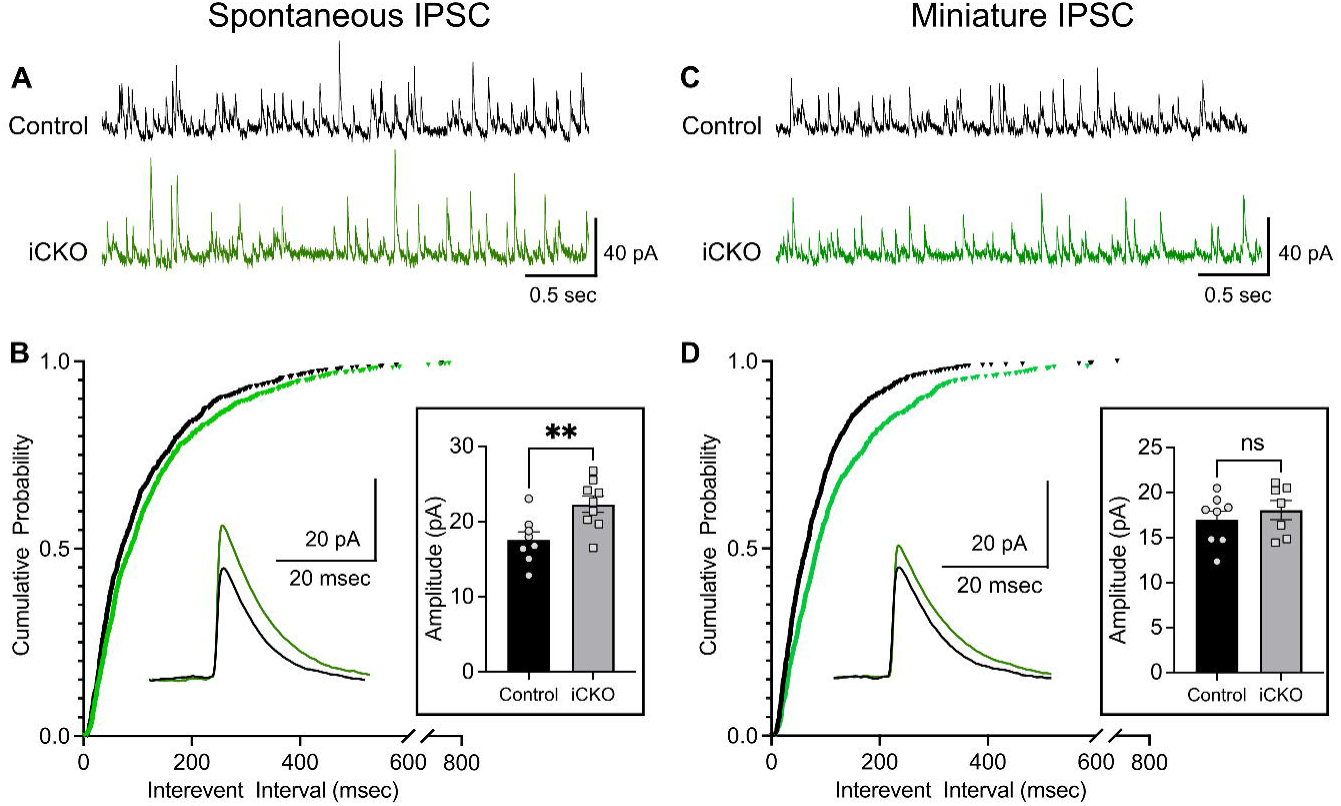
Reduced inhibitory synaptic inputs to CA1 PCs from iCKO mice. A) Representative spontaneous inhibitory postsynaptic current (sIPSC) recordings in control and iCKO mice. Note that events are less frequent in iCKO mice. B) Cumulative distribution of sIPSC interevent intervals show a right shift in iCKO mice suggesting a lower frequency of sIPSC compared to controls (n= 8-9 cells from 3 mice / group, p=0.0037, Kolmogorov-Smirnov test). Inset; the sIPSC amplitudes were larger in iCKO mice (p = 0.0071, unpaired t-test). C) Representative miniature inhibitory postsynaptic currents (mIPSC) in control and iCKO mice. D) Cumulative distribution of mIPSC interevent intervals show a right shift in iCKO mice indicating a lower frequency of mIPSC compared to controls (n= 7-8 cells from 3 mice/group, p<0.0001, Kolmogorov-Smirnov test). Inset; the mIPSC amplitudes were not different between control and iCKO mice. Representative averaged sIPSC/mIPSC traces are included for comparison.

Given our observation of reduced IPSC frequency, we investigated whether glutamatergic drive to CA1 PCs is altered in iCKO mice by evaluating genotype specific differences in sEPSC frequency and amplitude in CA1 PCs. CA1 PCs from iCKO mice showed an increase in the frequency of sEPSCs compared to controls. (Fig. 4A-B; sEPSC interevent intervals in ms: controls: 1424 ± 59.22, median: 765.7, IQR: 250.2 - 1872, *n*= 9 cells from 3 mice; iCKO: 1145 ± 44.28, median: 698.4, IQR: 265.7 – 1563, n= 9 cells from 3 mice; *p*= 0.0243 by Kolmogorov-Smirnov test, Cohen’s D: 0.17). However, sEPSC amplitude was not statistically different between groups (Fig 4B boxed inset; in pA, controls: 17.34 ± 1.127, iCKO: 17.29 ± 0.902, *n*=9 cells from 3 mice/group). Together, our findings demonstrate a deficit in basal inhibitory inputs and a maladaptive increase in excitatory inputs which could act in concert to enhance CA1 network excitability in iCKO mice.

**Figure 4:**
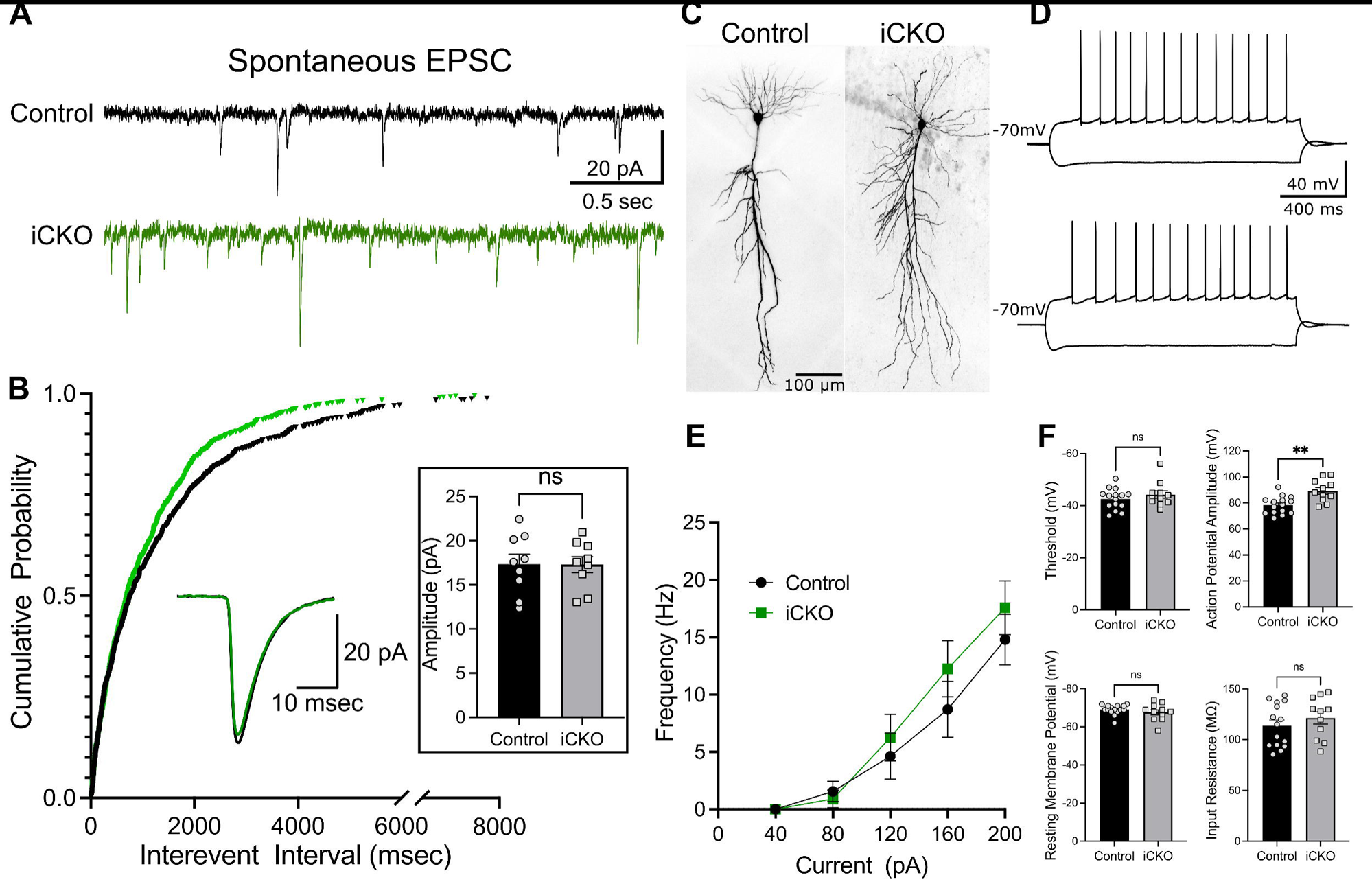
CA1 PCs from *iCKO* mice show changes in excitatory synaptic inputs but not intrinsic active and passive properties. A) Representative traces showing spontaneous excitatory postsynaptic currents (sEPSC) in control and iCKO mice. Note that events are more frequent in iCKO mice. B) Cumulative distribution of sEPSC interevent intervals show a left shift in iCKO mice suggesting a lower frequency of sIPSC compared to controls (n = 9 cells from 3 mice/group, p = 0.0243, Kolmogorov-Smirnov test). sIPSC amplitudes were not different between control and iCKO mice. C) Representative images of CA1 PCs filled during recordings. D) Membrane voltage response to hyperpolarizing (−200pA) and depolarizing (+200pA) step current injections from control (above) and iCKO mice (below). E) Summary plot of firing frequency in response to increasing current injections. F) Histograms compare firing threshold, resting membrane potential, input resistance and action potential amplitude in CA1 PCS between control and iCKO mice.

### *Nrp2* deletion in interneurons does not affect intrinsic physiology of CA1 PCs

Our findings show significant changes in inhibitory circuit function along with an increase in excitatory inputs in CA1 subfield after *Nrp2* deletion. Therefore, we evaluated whether the synaptic changes were accompanied by alterations in intrinsic active and passive properties of CA1 PCs. In current clamp recordings from CA1 PCs (Fig 4C) showed no differences in the frequency of action potential firing in response to step current injections (Fig. 4D-E). Although, firing threshold of CA1 PC’s were not different between control and iCKO mice, action potential amplitude was significantly increased (Fig 4F; Action potential threshold in mV: controls −42.58 ± 1.03, iCKO: −44.25 ± 1.44; amplitude in mV: controls 78.32 ± 1.72, iCKO 89.36 ± 2.55, *n* = controls 15 cells from 3 mice, iCKO 11 cells from 3 mice, *p*=0.0011, *t*_(24)_= 3.722 unpaired *t*-test). However, spike frequency adaptation and fast afterhyperpolarization (fAHP) in iCKO mice was not different from control (Supplementary Fig 3A-B, Spike frequency adaptation: controls 0.771 ± 0.086, iCKO: 0.582 ± 0.120; n = controls 15 cells from 3 mice, iCKO 11 cells from 3 mice). Examining passive membrane properties revealed no differences in resting membrane potential (RMP), input resistance (Rin), and sag ratio between controls and iCKO mice (Fig 4F and supplementary Fig. 3C. RMP in mV, control: −69.07 ± 0.658; iCKO: −67.64 ± 1.370; Rin in MΩ, control: 113.9 ± 5.415, iCKO: 121.5 ± 6.124; Sag Ratio: controls 0.955 ± 0.003, iCKO: 0.954 ± 0.003; *n* = controls 15 cells from 3 mice, iCKO 11 cells from 3 mice). These results demonstrate that CA1 PC active and passive intrinsic properties are mostly unchanged by selective deletion of *Nrp2* in interneurons.

### Selective deletion of *Nrp2* in interneurons increases risk for seizures

An imbalance in network excitation-inhibition ratio is considered a key factor in developing seizures. Our previous work on global Nrp2 deletion demonstrated an increased susceptibility to chemically induced seizure ^24^. However, whether selective deletion of *Nrp2* in MGE derived inhibitory precursors alone was sufficient to predispose the hippocampal network to seizures is not known. In addition, iCKO mice show a decrease in the number of inhibitory interneurons, a decrease in IPSCs and an increase in EPSCs suggesting altered excitation-inhibition balance. Unexpectedly, although the weight at the time of surgery and immediate post-surgical recovery did not differ between iCKO and control mice, iCKO mice had greater mortality 24-48 hours after electrode implant (50%, n=3 of 6) (Supplementary Fig. 4). Curiously, iCKO mice that died after surgery were found with extended hindlimbs suggesting that they may have had terminal seizures. Nevertheless, we did not observe spontaneous seizures in iCKO mice during handling, housing or during the 30 min baseline recording period. Following a single convulsive dose of KA (20mg/kg, i.p), iCKO mice showed a significantly shorter latency to convulsive seizures and reached status epilepticus more rapidly (Fig. 5A-B; latency in minutes, Controls: 27.7 ± 3.27, n=5; iCKO: 4.8 ± 1.95, *n*= 5 vs 3, *p*=0.0025, *t*_(6)_= 4.993 unpaired *t*-test). At 30 minutes post injection, all iCKO mice reached stage 4 seizures, classified using a modified Racine scale whereas, only 1 control mouse exhibited stage 4 seizure. Moreover, iCKO mice showed a 100% mortality within the first 60 minutes of KA induction whereas all control animals survived this period (Fig.5C; Mean latency to mortality in iCKO mice: 27.02 ± 12.22 minutes, p = 0.022, Mantel-Cox Test). Importantly, iCKO mice spent more time after KA injection (∼60%) exhibiting behavioral seizures compared to controls (Fig.5D, measured as total time spent in convulsive seizures/total time from induction to death or 60 minutes: Controls: 19.78 ± 4.42%; iCKO: 60.69 ± 3.12%, *n* = 5 vs 3, *p*<0.001, *t*_(6)_=6.483, unpaired *t*-test). Together, our experiments demonstrate iCKO mice exhibit greater seizure susceptibility than controls.

**Figure 5:**
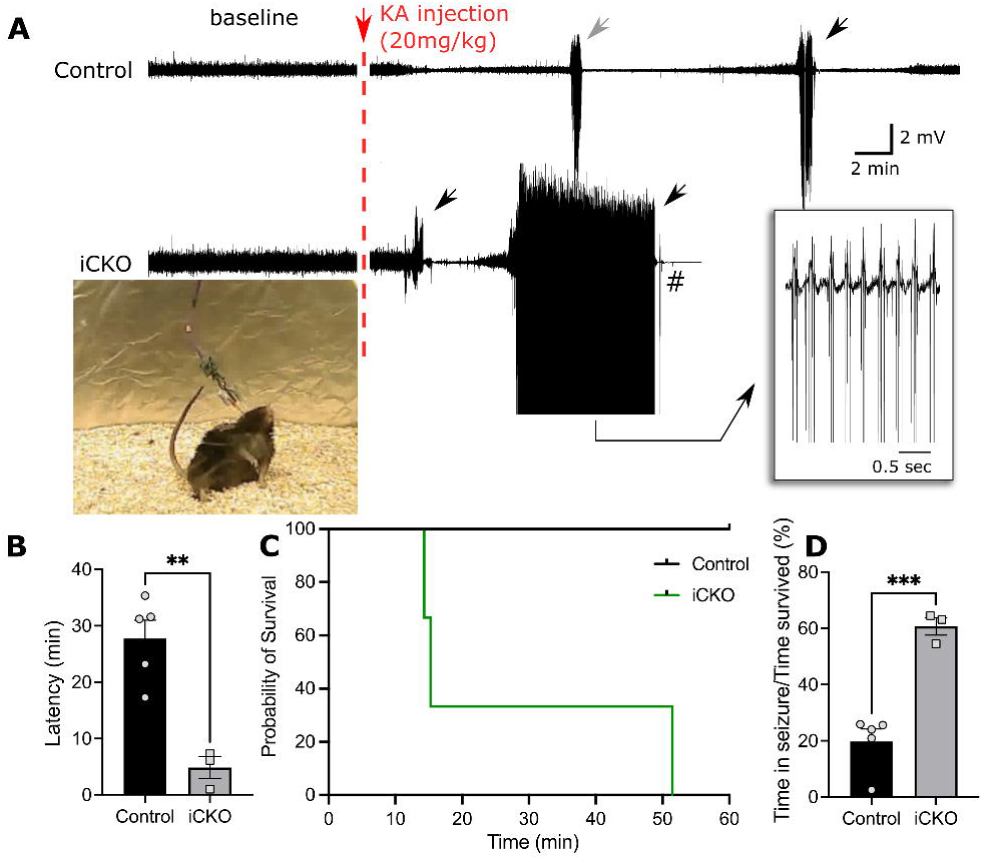
*Nrp2^f/f^;Nkx2.1-CreERT2+* mice exhibit higher susceptibility to KA induced seizures. A) Example in EEG traces show the baseline activity and the development of seizures following KA induction (20mg/kg). Note that iCKO mice have a shorter latency to seizure onset and have longer lasting seizures. B) Quantification of latency to first convulsive seizures after KA induction. Note that iCKO mice consistently had a short latency to convulsive seizures (n = 5 vs 3 mice, p = 0.0025, unpaired t-test). C) Survival analysis showing iCKO mice succumbed more often to seizures whereas controls animals survived the first 60 minutes after induction (n = 5 vs 3 mice, p = 0.0042, Log-rank Mantel-cox test.). D) Ratio between total time spend in seizures to time survived after KA injection shows iCKO mice spent 60% of their time after KA induction in seizure compared to 19% in control mice.

### Social novelty and goal directed behaviors are impaired in mice with interneuron targeted *Nrp2* deletion

Given *Nrp2*’s association with ASD in humans, and our previous results showing extensive social, learning, and sensorimotor alterations in *Nrp2*-null mice ^23^, we examined ASD-relevant behaviors in iCKO mice. We found that preference for social novelty (Fig. 6A), measured both as the proportion of time spent investigating (sniffing) the novel versus a familiar mouse (Fig 6B) and proportion of time mice spent within the chambers containing the novel and familiar mice (Fig 6C), was altered in iCKO mice compared to littermate controls. Unlike control mice, which showed a significant preference for the novel mouse, iCKO mice showed no preference for social novelty measured in the proportion of sniff duration directed at the novel and familiar mice (Fig 6B; *F*_(1,36)_=6.388, *p*=0.0110) or proportion of time spent within the chambers with enclosures housing the novel and familiar mice (Fig. 6C; two-way ANOVA interaction *F*_(1,36)_= 8.79, *p*=0.005, n=11 controls and 9 iCKO mice). As with the above measures, the ratio of the difference in time spent in the chamber with the novel versus familiar mouse to total time spent in the chambers with novel or familiar mice, a measure of social novelty preference showed and apparent decrease in iCKO mice, with the difference approaching statistical significance (Supplementary Fig 5A;, *p*=0.0504, unpaired t-test). Interestingly, evaluation of the time spent within the 3 chambers revealed that, in addition to spending more time in the chamber with the familiar mouse, iCKO spent significantly less time in the center chamber of the 3-chambered arena (Fig. 6D; *t*_(18)_ = 2.72, *p*=0.015, unpaired *t*-test). Further analysis failed to reveal genotype specific differences in overall sniff time (in sec, control = 110 ± 21 sec; iCKO = 94 ± 25 sec) indicating that the difference in social novelty preference is not due to an overall lack of interest in social interactions.

**Figure 6.**
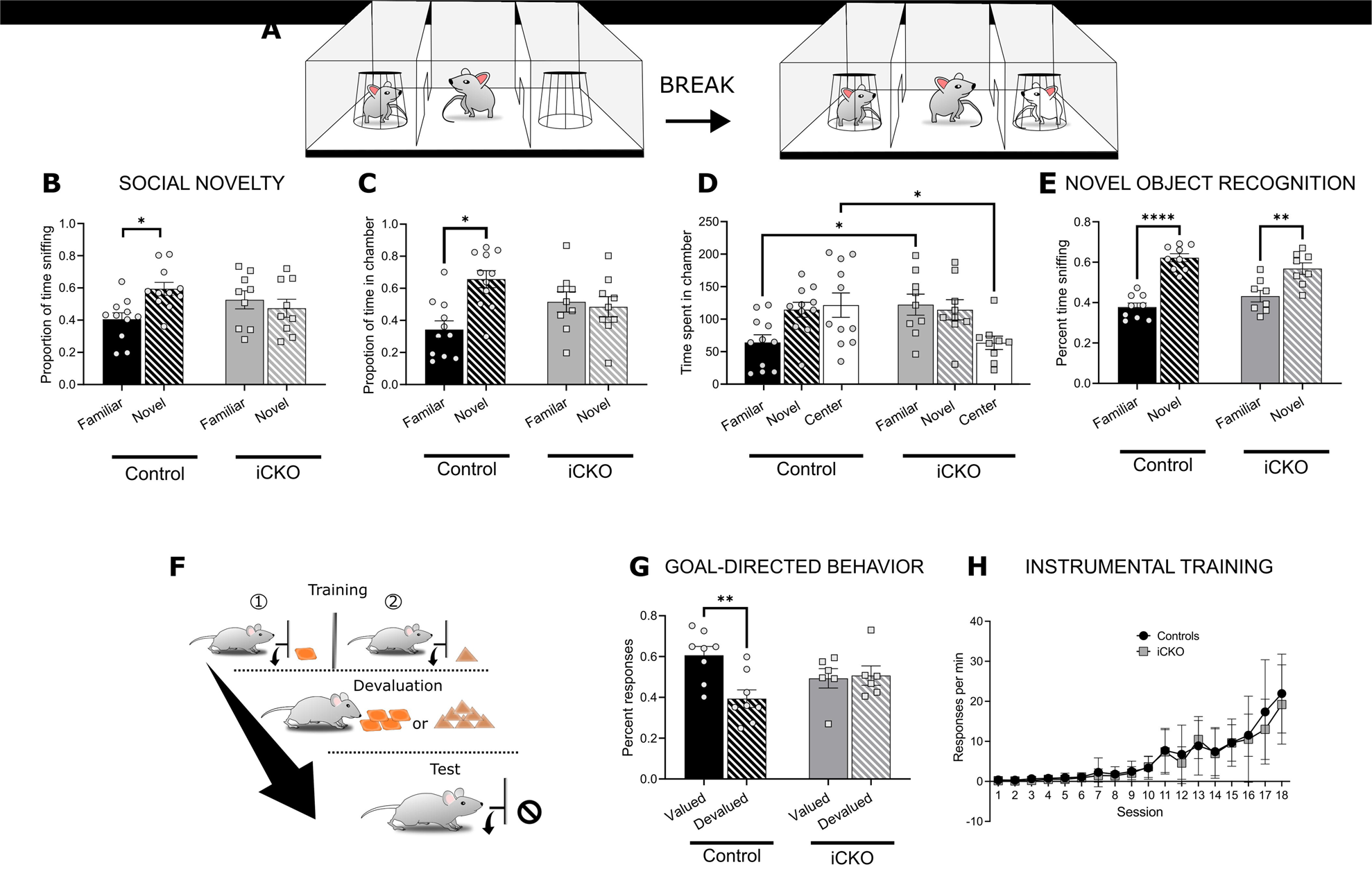
Social and goal directed behaviors are impaired in iCKO mice. A) Schematic of Social Novelty paradigm. B) Summary histogram compares the proportion of sniffs directed at novel and familiar mice. C-D) Histograms compare proportion of time spent within the chambers containing the novel and familiar mice (C), and time spent within each chamber during the social novelty test (D). Littermate controls spend a significantly greater proportion of their exploratory activity directed at a novel mouse compared to the lack of preference in iCKO mice (B: \**p*=0.011; C: \**p=*0.05; n=11 controls, n=9 iCKO). iCKO mice spend significantly less time in the center chamber compared to control mice (D: \**p*=0.015). E) Plot of time spent sniffing a novel compared to familiar object in a novel object task. Both littermate controls and iCKO mice spend more time exploring the novel object (\*\**p*=0.0011, \*\*\*\**p*<0.0001; n=9 controls, n=8 iCKO mice. F) Schematic of operant chamber training and devaluation. G) Histogram of percent responses to the valued and devalued outcomes in iCKO mice and littermate controls during a goal-directed task. Control mice made significantly more actions associated with valued outcomes, whereas iCKO mice showed no difference in selecting actions associated with valued and devalued outcomes (\**p*=0.02; n=8 controls, n=6 iCKO mice. Error bars are ± SEM. H) Summary data of response rate during training on the lever press task shows no statistical differences between the groups.

Since social preference test relies on memory of previous encounters, we tested whether deficits in recognition memory could have contributed to deficits in social novelty preference in iCKO mice. However, iCKO mice did not differ from control mice in an object recognition memory test (Fig 6E), indicating that the lack of preference for social novelty reflects a specific change in social behavior in iCKO mice.

In addition to changes in social behavior, iCKO mice differed from control mice in goal-directed behavior (Fig.6F). We trained mice to perform two different actions (right or left lever press) for two different types of food pellets (chocolate-flavored or grain pellets). Performance on this task provides evidence whether mice’s behavior is goal-directed or habitual. Evidence of goal-directedness is found when mice adjust their behavior when the value of the instrumental outcome changes. The selective-satiety procedure, in which animals consume one of the instrumental food outcomes to satiety, causes that outcome to lose value. Hence, when given the choice between actions for the devalued outcome or the valued outcome, goal-directed mice should preferentially respond on the lever for the still-valued outcome. In contrast, mice responding in a habitual manner should show no preference for the valued action compared to the devalued action.

We found the selective-satiety induced devaluation of one instrumental outcome significantly shifted the choice of the control mouse to the action (lever press) associated with the valued outcome, whereas the actions of iCKO mice showed no sensitivity to changes in outcome value (Fig. 6G; two-way ANOVA interaction *F*_(1,24)_=6.17, *p* = 0.02; *n*=8 controls and 6 iCKO). Importantly, iCKO mice and controls did not differ in acquisition of the instrumental response (Fig. 6H) demonstrating iCKO mice were capable of acquiring and performing a lever press action and showed no learning deficits. Similarly, locomotor measures in the open field and accelerating rotarod were not different between iCKO mice and controls (Supplementary Fig 5B-C) indicating lack of motor impairments and anxiety (distance traveled and time spent in center of arena p=0.81 and p=0.46 unpaired t-tests, respectively; rotarod p=0.53 two-way ANOVA). Moreover, control and iCKO mice did not differ in the time spent in open arm of the elevated-zero maze or grooming behavior (Supplementary Fig 5D-E, zero maze p=0.69 unpaired t-test; duration or frequency of grooming bouts p=0.93 and p=0.21, respectively unpaired t-tests). Therefore, the inability to adjust actions to changes in outcome value in iCKO mice strongly suggest a selective impairment in control of goal-directed actions.

Together, these findings suggest that Nrp2-dependent interneuron circuit development is a critical factor determining proper social behavior and cognitive flexibility in mice, and deficits in Nrp2 in inhibitory neurons during development can significantly impair these functions.

## Discussion

Dysregulated excitatory/inhibitory (E/I) balance is often considered a unifying pathology underlying ASD and epilepsy syndromes. Altered synaptic connectivity, loss of interneurons or hypoactive interneurons can contribute to shift in functional E/I balance ^8^. The emerging role of interneurons as critical determinants of circuit dysfunction in ASD is evident from studies in animal models and humans ^2,11^. Despite this knowledge, the molecular determinants of altered inhibitory circuits during neurodevelopment and the impact of such maladaptive circuits on network and behavioral function are unknown. Neuropilin-2 and its interactions with its ligand the secreted semaphorin 3F are crucial in establishing normal migration of interneuron to the cortex, the lack of which has been shown to result in increased NPY+ expressing interneurons in the striatum^20^. Importantly, Nrp2 is a strong candidate ASD gene on SFARI Gene database (https://gene.sfari.org/database/human-gene/NRP2#reports-tab). This is consistent with reports of polymorphisms in Nrp2 gene in patients with autistic syndromes in two distinct studies ^16,17^. We previously demonstrated that global Nrp2 deletion results in a significant decrease in density of several MGE-derived interneuron subtypes in hippocampus and reduced hippocampal inhibition ^24^. However, whether selective Nrp2 deletion in interneuron precursors, and not glutamatergic neurons, is sufficient to alter circuit function and contribute to ASD related behaviors is not known. The current study focused on the hippocampus to evaluate cellular and circuit effects of Nrp2 deletion in MGE-derived interneuron precursors. Our data demonstrate that, similar to the global *Nrp2* knockout ^24^, embryonic deletion of Nrp2 selectively in MGE-derived interneuron precursors leads to a significant reduction in PV+, SOM+ and NPY+ interneurons in the hippocampus (Supplementary Fig 6). While PV and SOM neuron counts were not different between mice with global *Nrp2* deletion (Supplementary Fig 6) and iCKO mice in the current study (mean cell count, PV: Global KO: 36.06±2.5, iCKO: 39.17±1.8, p=0.38; SOM: Global KO: 51.93±2.39; iCKO: 52.83±7.5, p=0.9), NPY neurons were lower in iCKO mice (mean cell count, Global KO: 67.98±3.46; iCKO: 47.83±1.93, p=0.003). These data demonstrate that interneuron specific embryonic deletion of Nrp2 is sufficient to reduce hippocampal PV, SOM and NPY neurons to the extent observed following global *Nrp2* deletion. Reduction in interneuron numbers in the hippocampus in mice with Nrp2 deletion could result from altered migration, integration or cell death, and result in altered interneuron numbers in other brain regions, such as the striatum or cortex. Alternatively, failure of MGE-derived precursors to reach their final destination could result in their elimination from the circuit by cell death. Although our analysis focused on the hippocampus, it is likely that embryonic *Nrp2* deletion in MGE-derived interneuron precursors has a broad impact on cortical interneuron numbers, as was observed following global *Nrp2* deletion (unpublished observations), which could impact behavioral outcomes.

At a circuit level, CA1 PCs from iCKO mice received a less frequent synaptic inhibition compared to control mice. This finding is similar to our observations in global *Nrp2* KO mice and could directly be attributed to the decrease in interneuron population in CA1^24^ (Supplementary Fig.6). The absence of a GABAergic inhibitory tone is believed to reduce signal to noise ratio and disrupt signal processing in individuals with ASD ^7,8,43^. We identify reductions in three major interneuronal classes in the iCKO mice; the soma-targeting fast-spiking PV+ basket cells and dendrite targeting SOM+ interneurons which are crucial regulators of neuronal spiking and integration of information ^44^, and NPY+ interneurons which are critical regulators of excitatory synaptic transmission and spiking frequency^45^. Therefore, the combined reduction in their populations is likely to have a cumulative effect on network function leading to alterations in excitatory synaptic transmission, neuronal oscillations, and theta/gamma coupling. Together, these changes could disrupt information processing and lead to behavioral deficits consistent with ASD ^46^. Coupled with the reduction in inhibition, we observed a significant increase in excitatory synaptic transmission in CA1 PCs from iCKO mice which could arise from an increase in circuit excitability due to loss of inhibition or from maladaptive plasticity. Moreover, while the deficits in inhibition in the iCKO mice are similar to those reported in the global *Nrp2* KO, the changes in CA1 PC excitability and resting membrane potential observed in the global *Nrp2* KO were not observed in the iCKO mice^24^ (Supplementary Fig.6). Thus, our findings support the proposal that interneuron specific knockout of *Nrp2* undermines E/I balance at a circuit level predominantly through disruption of the fine-tuned, layer-specific inhibitory control of CA1.

Enhanced excitability and deficient inhibitory control are fundamental mechanisms underlying seizures generation. Moreover, global *Nrp2* deletion leads to handling induced behavioral seizures at the time of weaning, ∼P21-P28 (unpublished observations) as reported previously ^47,48^. Similar to our prior findings in the global *Nrp2* KO, iCKO mice exhibited higher susceptibility to evoked seizures compared to age matched controls. Importantly, although our 30 minute baseline evaluation did not identify spontaneous seizures, the selective increase in postoperative mortality (Supplementary Fig. 4), as well as more severe and longer evoked seizures and higher mortality during evoked seizures in iCKO mice indicates their heightened seizure susceptibility. These findings are in line with prior reports in mice with global deletion of the high affinity ligand for Nrp2, Sema3F, which develop spontaneous seizures^49^. While we employed the power of mouse genetics to assess interneuron specific effects of the loss-of-function of Nrp2 on seizure susceptibility, gene polymorphisms are unlikely to be cell-type specific in clinical settings. Nevertheless, prior studies have noted that some molecular changes can differentially impact development of ASD-like behaviors and seizures ^50^ making it important to evaluate the contribution of interneuron-selective Nrp2 deletion to seizure susceptibility and neurobehavioral outcomes. Although our demonstration that seizure susceptibility is enhanced in *Nrp2* deleted mice is of potential clinical relevance, the two studies that identified Nrp2 polymorphisms in humans with ASD ^16,17^ explicitly exclude subjects with other neurological disorders making it difficult to assess seizures/epilepsy in their patient population.

Altered social behavior and impaired action control are core features of ASD. A diagnosis of ASD includes deficits in social communication or interaction, and restricted, repetitive behaviors^51^. Previously we showed that global *Nrp2* deletion results in broad impairments in motor coordination and repetitive movement, learning and memory, and mood and social behavioral tests (23,46). While our cellular analysis focused on the hippocampus, it is likely that embryonic *Nrp2* deletion in MGE-derived interneuron precursors has a broad impact on cortical circuits and impact behavioral function. In iCKO mice, we observed a lack of preference for social novelty which depends, in part, on hippocampal processing ^52,53^. iCKO mice engaged in social behavior, and indeed spent significantly less time in the center zone of the 3-chambered arena during the social novelty test. Despite engaging in social behavior, iCKO mice did not demonstrate a preference for social novelty that we observed on control mice. Instead, iCKO mice spent more time with the familiar mice without reduction in time spent with the novel mouse suggesting impaired learning or recognition of the familiar mouse. The reduced center time may reflect reduced fear-related or hyperlocomotive behavior, although we found no differences in elevate zero maze and open field behavior. Interestingly, unlike the global *Nrp2* knockout, iCKO mice showed no deficits in object recognition memory as assessed in the novel object recognition test (Supplementary Fig.6). These data indicate that the lack of preference for social novelty was not due to an impairment in episodic memory, or a lack of interest in novelty *per se.* The dependency of novel object recognition on hippocampus is debated, with more evidence in support of entorhinal cortex underlying performance in the novel object recognition task ^54^. We previously reported that global *Nrp2* KO animals were impaired in novel object recognition ^23^; however, global *Nrp2* KO also displayed increases cortical pyramidal neuron dendritic spine number and altered cortical and hippocampal pyramidal neuron excitability ^18,55^, which may have affected entorhinal cortex processing and produced a deficit in the novel object recognition task. Future studies should examine whether deficits in novel object recognition in the global *Nrp2* knockout result from altered Nrp2 regulation of glutamatergic cortical circuits. Regardless, our findings further underscore the functional distinctions between effects of global versus inhibitory neuron specific *Nrp2* deletion.

Goal-directed action was also impaired in iCKO mice. In an instrumental outcome devaluation task, mice are given the choice of actions associated with either a valued or devalued outcome. Unlike control mice that chose the action associated with the valued outcome, iCKO mice showed no preference. The impairment we observed in this task is unlikely to be caused by impaired sensory processing: we previously found that global *Nrp2* KO have intact olfactory discrimination and are motivated by food rewards ^23^. Furthermore, some KO mice responded at a higher rate for a particular pellet flavor during instrumental training, which would not occur if the pellets were indistinguishable. The impairment in goal-directed response in iCKO mice may be attributable to altered hippocampal activity. The dorsal hippocampus is transiently involved in the formation of action-outcome associations ^56,57^, which could have prevented mice from forming the associations necessary to guide their behavior in the devaluation test. Another possibility is that the altered interneuron migration induced in iCKO mice misplaced striatal interneurons ^20^. Disruption of striatal GABAergic interneurons impairs goal-directed behavior ^58,59^, suggesting striatal deficits as a potential source of impaired goal-directed behavior in iCKO mice. Future studies will address Nrp2’s role in striatal interneuron migration and its effects on behavior.

Certain limitations in generalizing the outcomes of our study to patient populations with Nrp2 polymorphisms are worth considering. This study was specifically developed to probe the impact of Nrp2 regulation of interneuron circuit development. In order to understand the developmental processes affected by Nrp2 deletion, we aligned timing of tamoxifen activity (administered at E12.5 and E13.5 and eliminated by P1) to the timeframe for interneuron migration and before programmed interneuron cell death, synapse development or plasticity (Supplementary Figure 1). Although the half-life of Nrp2 is unknown and could have impacted outcomes, our finding that iCKO mice have similar or fewer hippocampal PV, SOM and NPY neurons indicates that the timing of tamoxifen induction adopted here was appropriate to impact interneuron circuit development. While it is unlikely that human polymorphisms show temporal or cell type specificity, human Nrp2 polymorphisms are directly implicated in human ASD. However, a role for Nrp2 (or Sema3F) in human epilepsy is currently unknown. Despite our inability to assess spontaneous seizures due to high postoperative mortality, our results suggest that the altered circuitry resulting from Nrp2 deletion in interneuron precursors is sufficient to predispose the network to higher risk of seizures. Importantly, our results demonstrate that selective deletion of *Npr2* in interneuron precursors reproduces the inhibitory deficits observed in mice with global *Nrp2* deletion and specifically impairs ASD related behaviors.

Taken together, our findings provide a novel insight into the circuit and behavioral effects of embryonic *Nrp2* deletion in interneuron precursors. Our studies using interneuron specific *Nrp2* deletion combined with circuit and behavioral analyses support a role for developmental interneuronopathy in the co-occurrence of ASD related behaviors and risk for seizure disorders.

## Supporting information

Subramanian _Supplementary Methods

Subramanian_ Supplementary Figures

## Acknowledgements

This work is supported by the NJ Governor’s Council for Medical Research and Treatment of Autism: CAUT17BSP011 to V.S. and T.S.T., CAUT17BSP022 to T.S.T. and M.W.S., Rutgers Brain Health Initiative Pilot Grants to T.S.T. and V.S., NIH F31NS131052 and AES957615 to A.H., NIH R01 NS069861 and NS097750 to V.S., and NSF/IOS: 1556968 and 2034864 to T.S.T.

## Author contributions

D. S., C.E., A.H., J.B., and H.N. performed experiments; D. S., C.E., A.H., M.W.S., V.S., and T.S.T. analyzed data and interpreted results of experiments; D.S., C.E., and A. H. prepared figures; M.W.S., V.S., and T.S.T. conception and design of research; D.S, C.E., M.W.S., V.S. and T.S.T. drafted manuscript.

