## Supplementary material for "Dysregulation of Neuropilin-2 Expression in Inhibitory Neurons Impairs Hippocampal Circuit Development and Enhances Risk for Autism-Related Behaviors and Seizures": Subramanian _Supplementary Methods

**Animals**

All experiments were performed in accordance with IACUC protocols approved by Rutgers University, Newark, NJ, and the University of California at Riverside, CA and in keeping with the ARRIVE guidelines. The Neuropilin-2 (*Nrp2*) floxed conditional knockout mouse (*i*CKO), expression pattern was previously described (1). After Cre recombination, part of exon 1 is removed and replaced with tauGFP-pA+ sequence immediately downstream of the *Nrp2* promoter. Therefore, green fluorescent protein (GFP) is expressed in the neurons with successful *Nrp2* excision. The *Nrp2* floxed mouse was crossed with the *Nkx2.1-CreERT2* mouse (stock #014552), where Cre recombinase is induced to be expressed specifically in interneuron progenitors originating from the MGE (2). A total of 5 mg of tamoxifen was administered by oral gavage to the pregnant dams at E12.5 and E13.5, in accordance with interneuron migration peak timepoints. Cre recombinase was induced and excised *Nrp2* from the interneuron progenitors between approximately E12.5 and E14.5(3) in cells expressing the transcription factor Nkx2.1. Tamoxifen administration to the pregnant dam required delivery of the pups by C-section and placing pups with a wild-type foster mother. Deletion of *Nrp2* in the hippocampus was ascertained by visualization of GFP expression (Fig. 1). Mice used in these studies were backcrossed for at least 10 generations to the *C57BL/6NTac* background strain. Nrp2 genotypes were confirmed by using polymerase chain reaction (PCR).

**Immunocytochemistry**

For immunolabeling of tissues from mature adult *Nrp2f/f;NkxCre-ERT*2 mice: *Nrp2^+/f^;- or Nrp2^f/f^;-* littermate controls *and Nrp2^f/f^;NkxCre^+^* (iCKO) mice (average age of all genotype groups: 3.3 ± 0.7 months, four females and five males) (6). Mice underwent transcardial perfusion with 4% paraformaldehyde (PFA). Brains were dissected, postfixed for 2 h in 4% PFA, cryopreserved with 30% sucrose, and then imbedded in OCT medium. Brains were sectioned on a cryostat at 20μm thickness at 200μm intervals through the anterior to the posterior extent of the hippocampus along the coronal plane. Sections were processed for immunocytochemistry as previously described (7). Primary antibodies used were mouse monoclonal anti-Parvalbumin (1.5:500, Swant, PV235); mouse monoclonal anti-Parvalbumin (1:750, Sigma, P3088); rabbit polyclonal anti-Neuropeptide Y (1:1000, Abcam, ab30914); rat monoclonal anti-Somatostatin (1:150, Millipore, MAB354); chicken anti-GFP (1:750, Aves, GFP 1010), and for visualization, AlexaFluor 488 or Cy5 (1:500, Jackson Immuno Research Laboratories); AlexaFluor 546 (1:500 Invitrogen). After the final buffer wash, coverslips were mounted on microscope slides using Mowiol (Sigma Millipore, cat No. 81381), plus 10% p-Phenylenediamine (cat No. 78460) anti-fade mounting media.

**Cell counts and data analysis**

For cells containing parvalbumin (PV), neuropeptide Y (NPY), and somatostatin (SOM), the soma was counted in five major regions: dentate gyrus/hilar region/CA4 (DG) region, CA1, CA2, and CA3 regions. Subregions were defined by The Mouse Brain in Stereotaxic Coordinates (8) Demarcations were manually drawn on each section, as previously shown in Gant et al., 2009. (9) Interneurons falling fully within the demarcations were quantified using ImageJ software.

Within these regions, PV, NPY, and SOM-stained cells were quantified in the entire hemi-hippocampus. Ten PV, NPY, SOM, and DAPI stained hemi-hippocampal sections (at 200 μm intervals) per animal were counted using the ImageJ (Fiji) software. Experimenters, a total of two who independently counted the cells, were blinded to the genotypes. Data are represented as means per animal were used for statistical analysis (n = at least three animals/genotype). The total number of PV, NPY, and SOM cells were counted in the hippocampal region in each of 10 sections. Total counts were averaged together to establish a single value of cell counts per hemi-hippocampal region for each mouse.

**Photo-documentation**

All immunocytochemically processed brain sections were imaged using the Zen software (Zeiss) and Zeiss 510 LSM and Zeiss 980 LSM Confocal Systems. Representative nonoverlapping fields were imaged and stitched together to reconstruct hippocampal sections. Compiled z-stack images were exported as tif files. Images were captured with EC Plan-Neofluar 10X/0.30 M27, and Plan-Neofluar 40x/1.3 Oil DIC and the following laser lines: 405, 488, 543, and 633 nm.

**Ex vivo physiology**

Horizontal hippocampal slices (350µm) were prepared from 3 *Nrp2^+/f^;- or Nrp2^f/f^;-* littermate controls *and 3 Nrp2^f/f^;NkxCre^+^* (iCKO) mice (average age of all genotype groups: 12 months ± 3.49 days, four females and two males) following isoflurane anesthesia. Slices were prepared in ice-cold sucrose artificial cerebral spinal fluid (Sucrose-aCSF) containing (in mM): 85 NaCl, 75 sucrose, 24 NaHCO3, 25 glucose, 4 MgCl_2_, 2.5 KCl, 1.25 NaH_2_PO_4_, and 0.5 CaCl_2_. Slices were bisected and incubated at 32°C for 30 min in an interface holding chamber containing an equal volume of sucrose-aCSF and recording aCSF and were subsequently held at room temperature for an hour. The recording aCSF contained (in mM): 126 NaCl, 2.5 KCl, 2 CaCl_2_, 2 MgCl_2_, 1.25 NaH_2_PO_4_, 26 NaHCO_3_, and 10 D-glucose saturated with 95% O_2_ and 5% CO_2_ (pH 7.4). Whole-cell voltage-clamp recordings were performed at 33-34°C under IR-DIC visualization using MultiClamp 700B (Molecular Devices) as detailed previously (10,11).

Whole-cell voltage-clamp recordings of CA1 pyramidal cells (CA1-PCs) were performed at 33-34°C under IR-DIC using glass pipettes (3-6MΩ) fabricated from borosilicate glass capillaries using dual stage glass micropipette puller (Narishige PC-10, Amityville NY). CA1 pyramidal cells were patched under IR-DIC visualization using a Nikon Eclipse FN-1 microscope or Olympus BX50 with 40x water immersion objectives Whole cell voltage and current clamp recordings were obtained using MultiClamp 700B amplifiers, digitized at 10kHz using DigiData 1440A or DigiData 1550B and recorded using pClamp10 software (Molecular Devices, Sunnyvale, CA). K-gluconate based internal solution (in mM: K- gluconate 126, KCl 4, HEPES 10, Mg_2_ATP 4, Na_2_GTP 0.4, PO Creatine 10, pH 7.25, 270-285 mOsm) was used for recording intrinsic properties in current clamp mode. CA1 PCs were held at -70mV and voltage responses to 1500ms current steps were recorded (+200pA to -200pA at steps of 40pA). Active and passive properties included spike frequency, action potential amplitude, spike frequency adaptation ratio, resting membrane potential, input resistance, and sag ratio. Current clamp data was analyzed using Clampfit (Molecular devices, Sunnyvale CA). Sag ratio was calculated as the mean steady-state voltage from the last 100ms of the -200pA current step divided by the minimum voltage of the first 100ms of the -200pA current step. Input resistance was calculated by taking the mean steady-state voltage from the negative current steps (-200pA to 0pA) and taking the slope of the linear regression. Action potential detection was detected by Clampfit’s event detection. Threshold was determined by dV/dt > 0. Action potential amplitude was calculated as the height relative to the threshold. Spike frequency adaptation ratio was calculated as the ratio of peak-to-peak duration from the first two action potentials divided by the last two action potentials.

To record spontaneous and miniature excitatory/inhibitory currents from the same CA1-PCs, a cesium-based internal solution was used (in mM: Cesium methanesulfonate 140, NaCl 5, HEPES 10, EGTA 0.2, Mg_2_ATP 2, Na_2_GTP 0.2, Qx314 5, pH 7.25, 270-285 mOsm). Spontaneous and miniature inhibitory postsynaptic potentials (sIPSC and mIPSC respectively) were recorded at a holding potential of 0 mV, and spontaneous excitatory postsynaptic potentials (sEPSC) were recorded from the same neurons at a holding potential of -70 mV. Miniature currents were isolated by tetrodotoxin (TTX, 1μM). For analysis of interevent intervals and amplitudes of spontaneous/miniature synaptic currents, 100 events were detected using template search feature in Easy Electrophysiology (version 2.6) analyzed as cumulative distribution in Graphpad Prism 10.0.1

In vivo electrophysiology:

Eleven mice (5 *Nrp2^+/f^;- or Nrp2^f/f^;-* littermate controls) *and 6 Nrp2^f/f^;NkxCre^+^* (_iCKO); average age of all genotype groups: 9.12 ± 1.8 months, three females and eight males) were surgically implanted with a cortical screw electrode and a single tungsten wire depth electrode (50 μm, California Fine Wire company) in the CA1 subfield (AP:2 mm, ML: 1.5 mm, DV: 1.2mm from bregma). Two additional screw electrodes (invivo1, Roanoke, VA) on the contralateral side served as ground and reference. After 3–5 days of recovery, mice were connected to a tethered video-EEG monitoring system. Signals were sampled at 10 kHz, amplified (x100, 8202-SE3, Pinnacle Technologies), digitized (Powerlabs16/35, AD Instruments, Colorado Springs), and recorded using LabChart 8.0 (AD instruments). Following 30 minutes of baseline recordings, mice received a single high dose of KA (20 mg/kg), and their latency to electrographic seizure, seizure duration, and severity were measured. Seizures were scored by a blinded investigator based on a modified Racine scale as: Stage-1, absence-like immobility; Stage-2, hunching with facial or manual automatisms; Stage-3, rearing with facial or manual automatisms and forelimb clonus; Stage-4, repeated rearing with continuous forelimb clonus and falling; and Stage-5, generalized tonic-clonic convulsions with lateral recumbence or jumping and wild running followed by generalized convulsions(13) . Seizure severity scores in the first 30 min were averaged over 5 min epochs.

**Behavior testing**

*Instrumental goal directed behavior test*

Fifteen mice (7 *Nrp2^+/f^;- or Nrp2^f/f^;-* or *Nrp2^+/f^;NkxCre^+^* and 8 *Nrp2^f/f^;NkxCre^+^*); average age of all genotype groups: 5.2 ± 0.2 months were evaluated for decision making ability in this test. Mice were placed on a restricted food diet of approximately 2 g of standard chow each day. The chow (Purina, St. Louis MO, USA) was given in their home cage after behavioral procedures were complete. Animals were weighed daily, and their body weights were maintained to 85–90% of their original weight. Mice were tested in eight operant conditioning chambers (Med Associates, St. Albans VT, USA) as previously described(7). Operant chamber operation and data collection were carried out with Med Associates proprietary software (Med-PC V). While placed on food restriction, mice were simultaneously habituated to the operant conditioning chamber for one 15-min session. The next day, mice were conditioned to find pellets in the food cup in a 20-minute session in which food pellets were dispensed on a random-time 60 s schedule. Levers were retracted during this phase. The next day mice were trained to use levers.

Mice are first trained to press levers and associate levers with a given food pellet in operant conditioning boxes. Boxes are 15.9cm x 14.0 cm x 12.7 cm (w/h/d) and include a food cup and retractable levers which deliver 20mg grain-based chocolate flavored or grain flavored food pellets into the food cup. For the devaluation test, mice are placed in cages identical to their home cages. They are provided a bowl with 10g of one flavor of food pellet. After 1 hour, the mouse is placed in an operant conditioning box for 10 minutes and allowed to press one randomized lever associated with either the food item eaten or another food item. The following day, the test is repeated with 10g of the other type of food pellet provided in the bowl. (14). This devaluation test is operated, and data collected with Med Associates proprietary software (Med-PC IV).

*Novel object recognition test*

The ability to recognize and prefer a novel object over a familiar one is assessed. The mouse is allowed to explore two identical toys in a 40 cm X 40 cm open field arena for 10 minutes (Stoelting Co, USA). Toys are placed in opposite corners 10 cm from the walls. Then, mice are placed in their home cage for 30 minutes. A novel toy then replaces one of the familiar toys and the mouse is allowed to explore for 5 more minutes. The time and frequency spent sniffing the novel and familiar toys is assessed from video footage and quantified as a percentage of time spent sniffing both the novel and familiar toys. Noldus ethovision software is used to capture video and track the position of the mouse.

*Social novelty test*

The ability to recognize and prefer a novel mouse versus a familiar one is assessed. The test mouse is placed in the middle of a three-chambered arena separated by plexiglass barriers which have small holes allowing the mouse to pass. The entire arena measures 50 cm wide, 30 cm deep and has 20 cm high walls with each chamber measuring approximately 16 x 30 cm. The enclosure housing the novel/familiar mice has a 7 cm base diameter. Mice are habituated to the arena for 30 minutes. In the social preference phase, a caged mouse is introduced into one of the outside chambers and an empty cage to the other and the test mouse is allowed to roam for 10 minutes. During the social novelty 10-minute test phase, the caged mouse is now the “familiar” mouse and a new mouse is placed in the empty cage as the “novel” mouse. The location of the novel and familiar mice is randomized. The time and frequency spent sniffing the novel and familiar mouse or empty cage is assessed from video footage and quantified as a percentage of time spent sniffing both the novel and familiar mice. Noldus ethovision software is used to capture video and track the position of the mouse. Sniffs directed at the novel and familiar mice were scored when the test mouse’s nose was adjacent to the cage containing the stimulus mouse and the mouse adopted a sniffing posture. Mice were fully enclosed in the chamber containing the novel or familiar mouse when engaging in sniffing behavior.

*Accelerating rotarod test*

An accelerating rotarod for mice (Harvard Apparatus, Holliston MA) was used to test motor coordination. Mice were placed on the spindle, which began rotating at 4 rpm and linearly accelerated over 2 min to a maximum of 40 r.p.m. Latency to fall was recorded. If mice made a single revolution while gripping the rod the trial ended. Over two days, mice completed 5 trials per day with 10 min break between trials.

*Open field test*

Four standard mouse open field chambers (Stoelting, Wood Dale IL) were used to assess motor and anxiety behaviors. Mice were placed in the center of the chamber and removed after 20 minutes. A total of two trials were conducted, once per day over 2 days. An overhead camera (Basler Ace) recorded mouse movement. Boxes were illuminated with overhead LED panel lights at approximately 250 lumens. Total distance traveled and time spent in the center (20 cm x 20 cm) were calculated using the Noldus Ethovision software.

*Grooming test*

Self-grooming behavior was assessed in mice in a cage identical to their home cage for 20 minutes. Observers scored the duration and frequency of licking and scratching of their face or other body areas from video using the Noldus Ethovision software.

*Elevated zero maze*

Movement in an elevated zero maze (Stoelting) was assessed. Mice were placed on a circular track 50 cm above the floor. The circular track was 50 cm in diameter and had two areas that were enclosed by walls 15 cm in height, which total 25 cm in diameter. After 5 minutes, mice were removed from the track. Noldus Ethovision software was used to capture video and track the position of the mouse. Time spent in the open versus closed arms was assessed.

**Statistical analysis**

Unpaired *t*-tests, two-way ANOVA and post hoc Bonferroni’s, Sidak’s or Tukey’s multiple comparison correction (Graph Pad Prism Software) were used for statistical differences in the behavioral tests and cell count. The difference between two dependent means (matched pairs) was used to determine the sample size requirement of behavioral tests using G*Power 3.1 software. A sample size requirement of three to seven animals for cell counts and 5 to 8 for behavior testing was estimated by using 80% power and an effect size found in previous work and literature. Exclusion criteria of three standard deviations from the mean was pre-established for behavior tests. One mutant mouse performed over three standard deviations lower than the mean and was therefore eliminated from behavior analysis. Significance was set to p < 0.05. Data are shown as mean ± SEM.
