## Supplementary material for "Dysregulation of Neuropilin-2 Expression in Inhibitory Neurons Impairs Hippocampal Circuit Development and Enhances Risk for Autism-Related Behaviors and Seizures": Subramanian_ Supplementary Figures

A

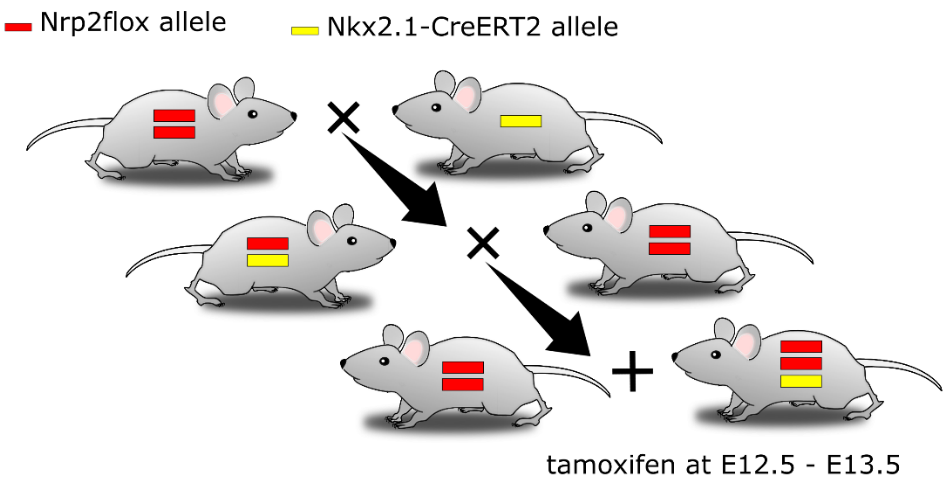


B

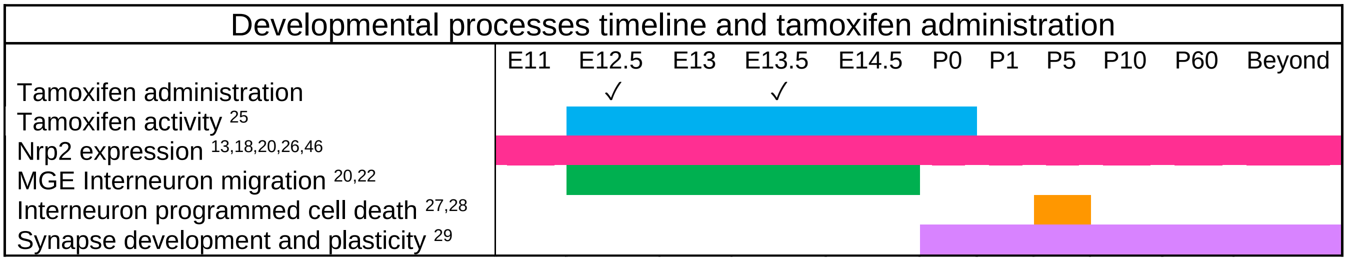


References correspond to bibliography in main text.

**Figure S1. Schematics of experimental model development.**

A. Schematic illustrates the inducible conditional knockout strategy which allows Nrp2 to be excised from neurons harboring transcription factor Nkx2.1 found in inhibitory neurons originating from the MGE in a spatiotemporal controlled manner. Nrp2flox mice were crossed with Nkx2.1-CreERT2 mice to generate mice with and without Nkx2.1-Cre positive alleles. B. Schematic of the timeline of interneuron developmental processes and tamoxifen administration targeting interneuron migration from MGE.

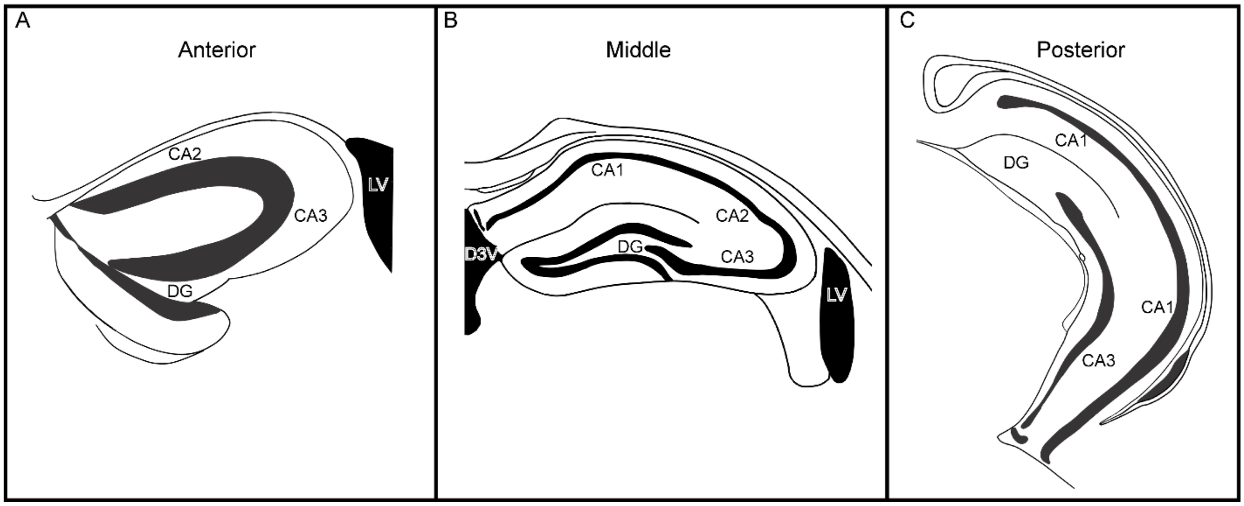


**Figure S2. Illustration of regions examined for cell counts across the septo-temporal axis of the hippocampus.**

A-C. Illustrations showing slices used for quantification of interneurons (A), anterior (B) middle (C) posterior hippocampal slices.

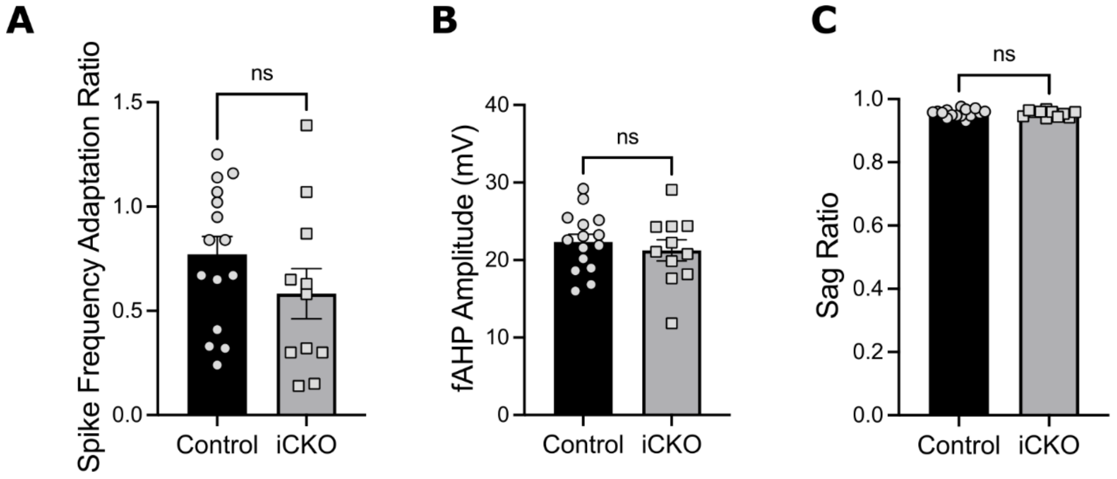


**Figure S3. CA1 PC intrinsic properties are not different between control and iCKO mice.**

A-C. Spike frequency adaptation ratio (A), fast afterhyperpolarization (fAHP) (B) and sag ratio (C) are not different between control and iCKO mice.

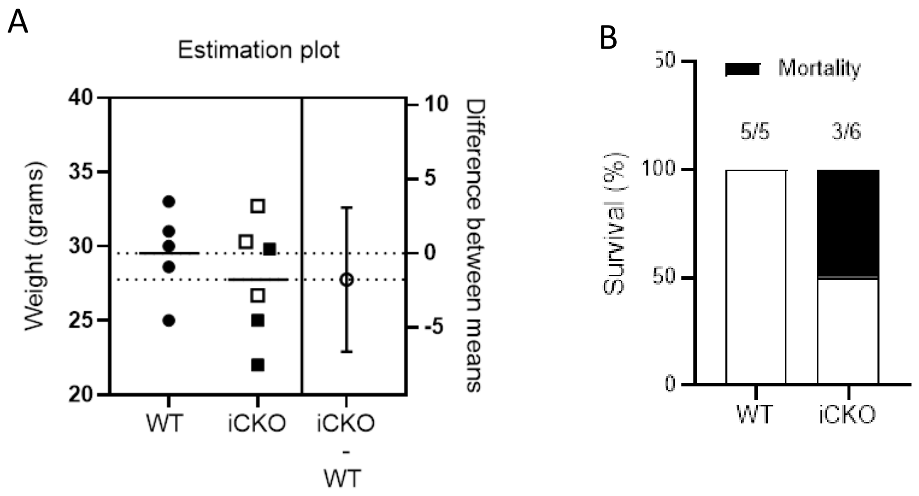


**Figure S4. Selective increase in postoperative mortality in iCKO mice.**

(A) Estimation plot shows no difference in the weight of the animals at the time of surgery (P=0.430; by Unpaired t-test, n = 5 WT and 6 iCKO). Note that open squares represent animals that died after surgery. (B). Plot of postoperative survival 48 hours after electrode implant shows 50% mortality in iCKO mice and no mortality in littermate controls.

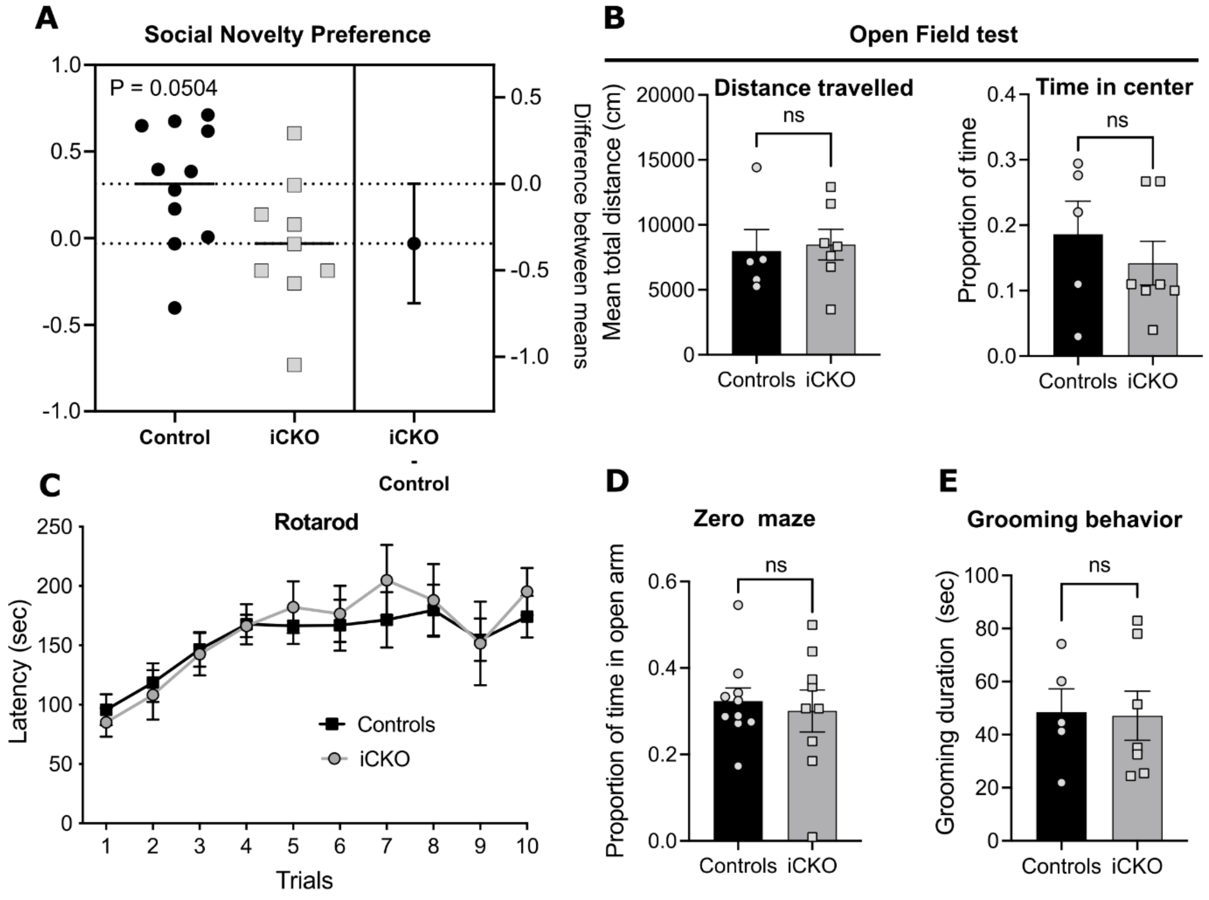


**Figure S5. iCKO mice show normal motor and anxiety behavior.** A. Estimation plot of social novelty preference calculated as the difference between time spent in chamber with novel mouse versus familiar mouse divided by total time spent in chambers with novel and familiar mice. B. Summary plots of total distance traveled and time spent in the center measured during an open field tests reveal no significant difference in performance between control and iCKO mice. C. Plot of latency to fall from the accelerating rotarod in the rotarod test of sensorimotor learning revealed no difference between groups. D. Summary of time spent in open arms of an elevated zero maze showed no differences between groups. E. Plot of the duration of grooming during a 20-min observation period was not different between iCKO and control mice (n=5 controls, n=7 iCKO mice).

| | Global Nrp2 KO | Interneuron-specific Nrp2 KO |
|---|---|---|
| **Hippocampal cell counts** | | |
| PV interneuron | ↓ | ↓ |
| NPY interneuron | ↓ | ↓ |
| SOM interneuron | ↓ | ↓ |
| **Physiology** | | |
| CA1 PC Intrinsic properties | ↓ | ⟺ |
| IPSC frequency | ↓ | ↓ |
| **Seizure susceptibility** (latency to convulsive seizure) | ↑ | ↑ |
| **Behavioral assay** | | |
| Grooming | ↑ | ⟺ |
| Rotarod | ↓ | ⟺ |
| Open Field | ⟺ | ⟺ |
| Novel object recognition | ↓ | ⟺ |
| Goal directed behavior | ↓ | ↓ |
| Social novelty | ↓ | ↓ |
| Zero maze [55] | ⟺ | ⟺ |

**Figure S6. Table comparing the of effects of Global vs Interneuron selective Nrp2 Knockout.**

Table showing cells counts, physiology and seizure susceptibility are altered in both global and interneuron specific Nrp2 KO compared to their respective WT controls. However, in contrast to global Nrp2 KO, iCKO mice showed no difference in behaviors related to grooming, rotarod, open field, zero maze and novel object recognition. Data for global Nrp2 KO is based on Eisenberg et al., 2021, Ref #24 and Assous et al Ref#55.
